# Profile of the *in silico* secretome of the palm dieback pathogen, a fungus that puts natural oases at risk

**DOI:** 10.1101/2021.11.18.469146

**Authors:** Maryam Rafiqi, Lukas Jelonek, Aliou M. Diouf, AbdouLahat Mbaye, Martijn Rep, Alhousseine Diarra

**Affiliations:** Institute of Bioinformatics and Systems Biology, Justus Liebig University, Giessen, Germany; Plant Pathology Program, Agrobiosciences, University Mohammed VI Polytechnic (UM6P), Benguerir, Morocco; Swammerdam Institute for Life Sciences, Faculty of Science, University of Amsterdam, The Netherlands; Digital 4 Research Labs, Mohammed VI Polytechnic University (UM6P), Benguerir, Morocco

## Abstract

Understanding biotic changes that occur alongside climate change constitute a research priority of global significance. Here, we address a plant pathogen that poses a serious threat to life on natural oases, where climate change is already taking a toll and severely impacting human subsistence. *Fusarium oxysporum* f. sp. *albedinis* is a pathogen that causes dieback disease on date palms, a tree that provides several critical ecosystem services in natural oases; and consequently, of major importance in this vulnerable habitat. Here, we assess the current state of global pathogen spread, we annotate the genome of a sequenced pathogen strain isolated from the native range and we analyse its *in silico* secretome. The palm dieback pathogen secretes a large arsenal of effector candidates including a variety of toxins, a distinguished profile of secreted in xylem proteins (SIX) as well as an expanded protein family with an N-terminal conserved motif [SG]PC[KR]P that could be involved in interactions with host membranes. Using agrobiodiversity as a strategy to decrease pathogen infectivity, while providing short term resilient solutions, seems to be widely overcome by the pathogen. Hence, the urgent need for future mechanistic research on the palm dieback disease and a better understanding of pathogen genetic diversity.

## Main

Global climate has changed rapidly over recent decades, and climate-change predictions in some water-limited regions, such as westernmost Mediterranean (Iberia and Morocco), North Africa and Middle East, forecast a significant shift in the near future, with less frequent precipitation and hotter and longer drought events (Varela et al., 2020, Weniger et al., 2019). An impact of climate change has already been observed on biodiversity and crop productivity; and consequently, on human livelihoods in affected areas (Wheeler & von Braun, 2013). Because of its desert location, vulnerable ecosystem and farming practices, oases are at major risk (Mohamed et al., 2021).

Alongside abiotic stresses caused by global warming, natural oases are increasingly facing threats by emerging infectious plant diseases, notably those caused by fungal and oomycete pathogens (Table 1). In terms of yield losses, fungi are the most destructive emerging pathogens in oases, posing a serious threat to food security and oasitic ecosystem health. While many emerging diseases in oasis ecosystems are encountered in other agroecological zones (Fones et al., 2020), a few diseases are exclusively of oasitic origin. One such disease is dieback of palm, also known as Bayoud, caused by the ascomycete fungus *Fusarium oxysporum* f. sp. *albedinis*. The palm dieback pathogen has been identified in the late 19^th^ century in the oasis of Zagora, Southern Morocco (Louvet & Toutain, 1973), but it has only started to spread rapidly in recent decades. Palm dieback has now been detected in many palm-growing areas across Africa, Asia and America (Sedra, 2013, Corte, 1973, Feather, 1979, Ibrahim Elkhalil Benzohra, 2015, Mirza Hussain Samo, 2020), with North Africa being the most affected. An estimated 15 million palm trees in oasitic habitats of Morocco and Algeria have now been completely wiped out by the palm dieback disease, which continues to progress at an alarming rate despite prophylactic measures to contain pathogen spread (Mirza Hussain Samo, 2020). When outbreaks of palm dieback take a pandemic scale, natural habitats collapse entirely (Figure1), severely impacting human livelihoods, economy and subsistence in populated oases.

**Table 1:** Emerging infectious plant diseases in natural oases.

| Species | Disease | Pathogen | References |
| --- | --- | --- | --- |
| Phoenix dactylifera | Bayoud disease | Fusarium oxysporum f.sp. Albedinis | (Toutain, 1965; Louvet and Toutain 1973 ; Sedra, 1992) |
| Phoenix dactylifera | Brittle leaf disease | Unknown | (Namsi et al., 2007) |
| Phoenix dactylifera | Lethal yellowing disease | Candidatus Phytoplasma palmae | (Harrison et al., 2008) |
| Phoenix dactylifera | Inflorescence rot disease | Mauginiella scaettae | (Taxana and Larous, 2003) |
| Phoenix dactylifera | Diplodia leaf-base disease | Diplodia phoenicum | (El-Deeb et al., 2007) |
| Phoenix dactylifera | Belaat disease | Phytophthora palmivora | (Sedra, 2003) |
| Phoenix dactylifera | Black scorch disease | Thielaviopsis punctulata | ( Al-Naemi et al., 2014) |
| Phoenix dactylifera | Bending Head Disease | Botryodiplodia theobromae | (Sedra, 2003) |
| Citrus aurantifolia | Witches' Broom disease | Candidatus phytoplasma aurantifolia | (Khan & Grosser, 2004) |
| Olea europaea spp. | Root rot disease | Cylindrocarpon destructans, Phytophthora spp. | (Zizzerini and Marte, 1976 ; SIPAM, 2008) |
| Malus pumila | Fire Blight disease | Erwinia amylovora | (Fatmi et al., 2008 ; SIPAM, 2008) |
| Solanum lycopersicum | Tomato wilt | Fusarium oxysporum f. sp. Lycopersici | (SIPAM, 2008 ; Murugan et al., 2020) |
| Solanum melongena | Root-knot disease | Meloidogyne arenaria | (SIPAM, 2008 ; Mokriani et al., 2019) |
| Vitis vinifera | Pierce's disease | Xylella fastidiosa | (Choi et al., 2013) |

**Figure 1.**
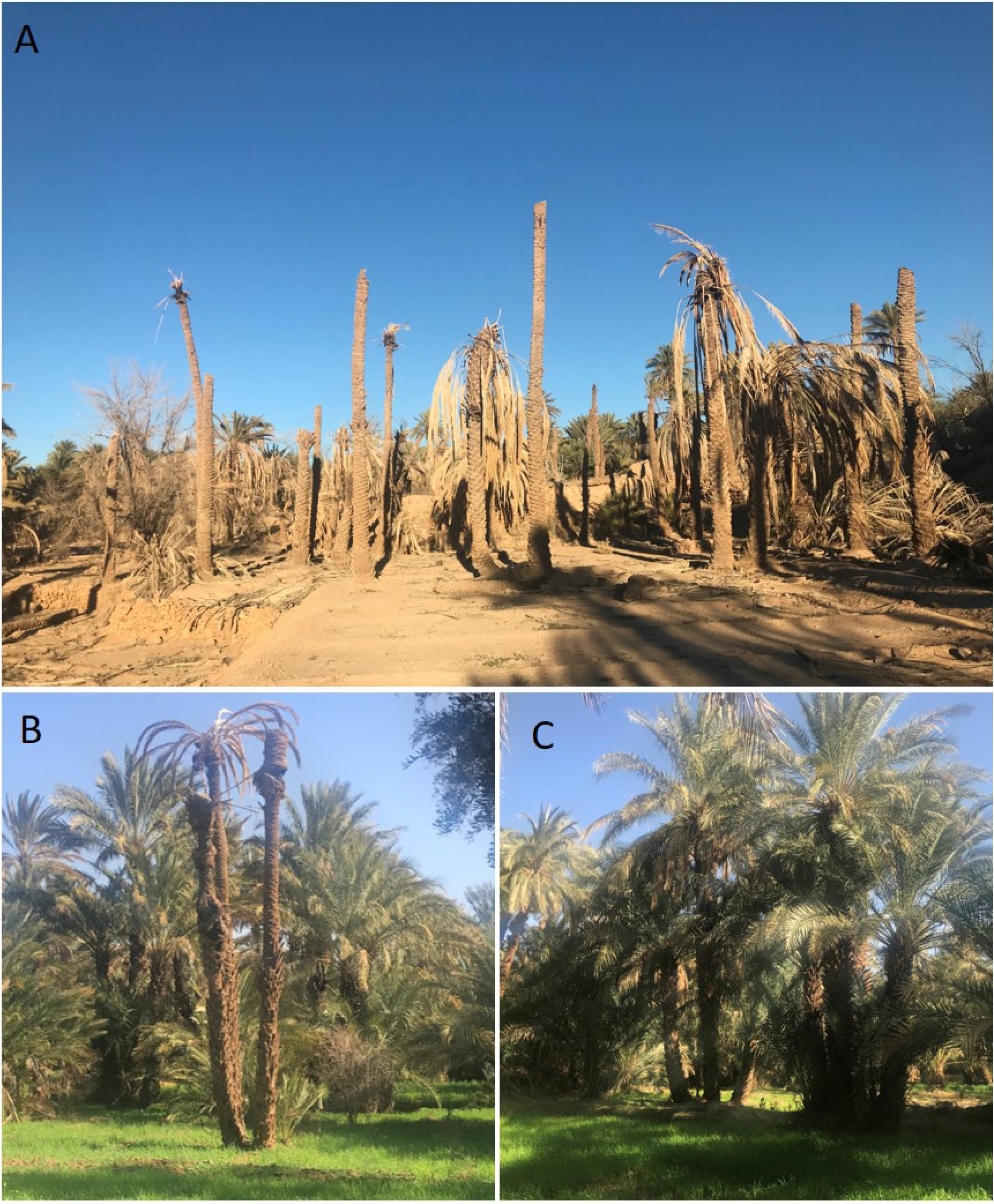
Dieback of date palm in a natural oasis in Morocco. (a) Date palm tree showing dieback on the crown. (b) Healthy date palm. (c) Palm dieback disease taking a pandemic scale and leading to ecological collapse of a Moroccan oasis

Date palm (*Phoenix dactylifera*) is a distinct species that belong to the palm family (Arecaceae of the monocotyledon order Arecales) (ros-Balthazard M., 2020a). Date Palms are pivotal components of natural oasis ecosystems, where they provide many ecosystem services, such as protection from silting and desertification, cooling oasis temperature, creating a suitable microclimate that supports the development of other crops, changing soil-water dynamics, and supporting wildlife as well as providing food and animal feed. Through these diversified ecosystem services, date palm contributes to the long-term functioning and, notably, the resilience of oasitic ecosystems to natural and anthropogenic perturbations (de Grenade, 2013). Most date palm cultivars are susceptible to the dieback disease and have already succumbed in North Africa despite efforts of restoration (Sedra, 2013, Ibrahim Elkhalil Benzohra, 2015). Palm dieback disease is jeopardising the genetic diversity of date palm in affected oases, especially when spreading at an epidemiological rate, wiping out groves and, sometimes, entire oases (Figure 1A). Besides, the number of palm trees lost to dieback disease may be underestimated, as farmers stop irrigating entire groves and regions of the oases where the disease has erupted as an effort to limit pathogen spread, thereby killing not only domesticated but also date palm’s wild relatives that represent a reservoir of genetic diversity, which is needed for agricultural improvement.

Palm dieback disease is characterised by discolouration from green to whitish of one palm leaf of the middle crown, followed by necrosis on the dorsal side of the rachis that progresses from the base to the tip of the frond, corresponding to mycelia progression in the vascular bundles of the rachis. After one leaf has been infected, adjacent leaves show the same succession of symptoms. This leads to crown dieback and ultimately the death of infected trees (Zaid, 2002) (Figure B). All commercialised date palm cultivars seem to be equally susceptible to the disease, which affects young offshoots as well as over 200-year-old palms. Albeit alarmingly destructive, pathogenesis of the disease is as yet uninvestigated and molecular details of how *F. oxysporum* f. sp. *albedinis* brings about infection of palm trees are still unknown. Secretome profiling and predictive ranking of effector candidates of this pathogen have, so far, not been addressed in research. It is therefore timely to address the palm dieback pathosystem in order to understand how the disease takes place and, subsequently develop disease control strategies that can save date palm trees and natural oases. Here, we annotate the draft genome of Foa strain 133 (Khayi et al., 2020), we mine its secretome, we analyse its large repertoire of effector candidates, and we briefly discuss research areas that should be the focus of future studies in order to get insights into Foa’s molecular mechanisms of pathogenicity.

## Results and discussion

### Annotation and secretome profiling of *F. oxysporum* f. sp. *albedinis*

To mine the secretome of the palm dieback-causing agent *F. oxysporum* f. sp. *albedinis* (Foa), we annotated the draft genome of the Foa 133 strain (Khayi et al., 2020), as described in the methods section. Annotation of the identified 16887 genes is summarised in supplemental data1. The secretome was determined by processing predicted protein datasets through an in-silico secretion pipeline (Figure 2). In this study, the secretome is defined as proteins that are predicted to be secreted extracellularly through the classical ER secretory pathway, do not target mitochondria and do not integrate into the membrane through transmembrane domains (TMs). Of the predicted 16887 genes, 1464 were predicted to code for secreted proteins, of which 1077 (73%) contained less than 500 amino acids and 598 proteins had no identified pfam domain (Table 2). Annotation of all secreted proteins is presented in table S1. Foa secretome harbours a large arsenal, 386, of carbohydrate active enzymes (CAZymes) that are predicted to be active in the apoplast, contributing mainly to plant cell wall degradation. These predicted CAZymes are distributed among glycoside hydrolases (GH), auxiliary activities (AA), carbohydrate esterases (CE), pectin lyases (PL), carbohydrate binding modules (CBM) and glycosyl transferases (GT) families, harbouring 209, 107, 26, 37, 9 and 3 members, respectively. 73 GH and CE proteins are accompanied by CBM modules (table S2). 550 predicted secreted proteins contain cysteines that are predicted to form bisulfide bridges. Of these, 437 proteins contain at least two predicted disulphide bonds (table S3). 313 secreted proteins were predicted to carry a wide range of enzymatic functions. 370 proteins (25%) contain repeats, carrying two or more copies of a tandemly or non-tandemly duplicated sequence or structural motif that is at least five amino acid residues in length. We have not considered degenerate sequence repeats that may be identifiable only through analysis of protein tertiary structure. Proteins carrying LysM domains were also recovered in the secretome. One protein (FUN_010192-T1) carried *Alternaria alternata* allergen 1 pfam domain PF16541.4.

**Figure 2.**
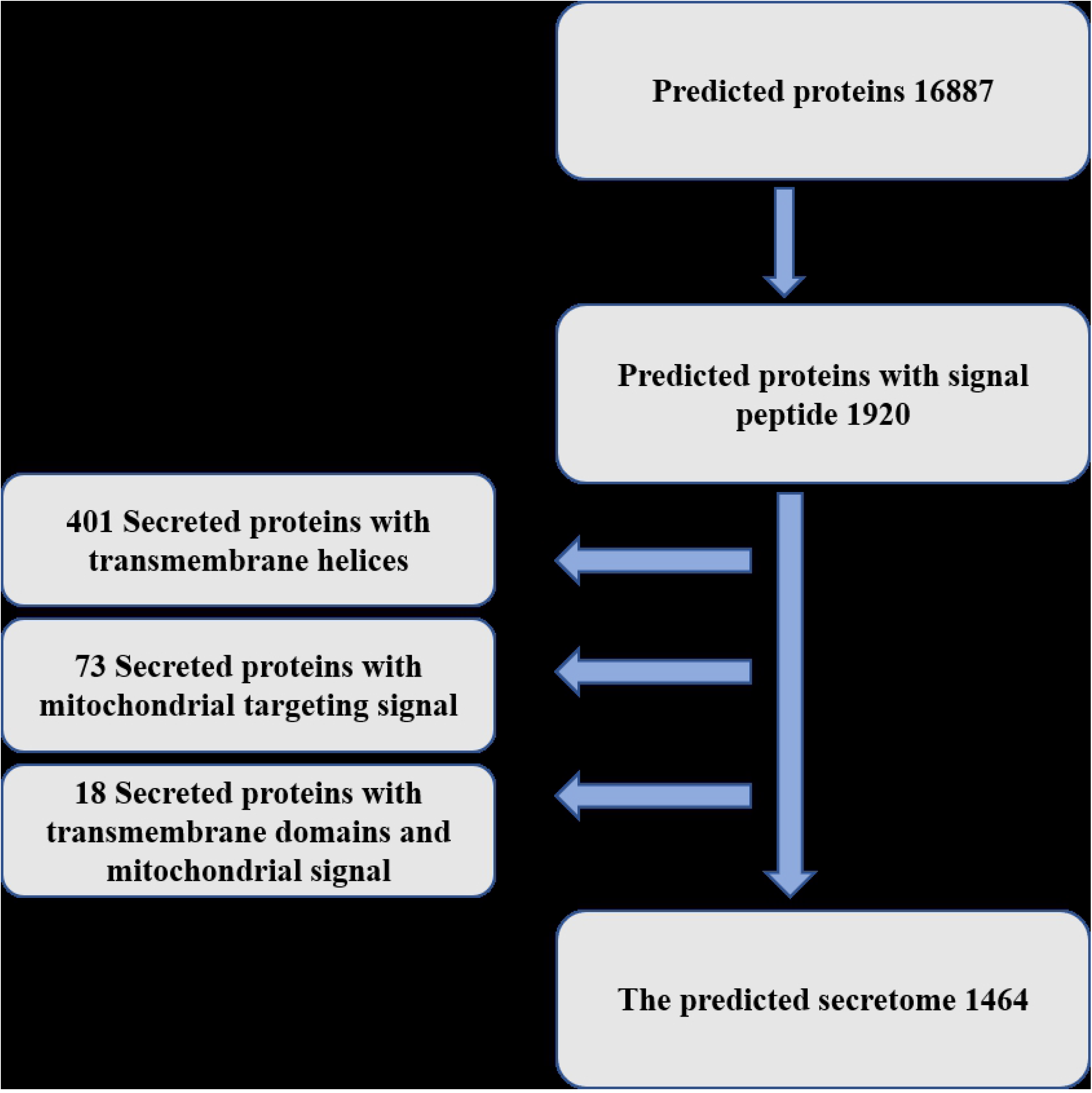
Overview of the computational pipeline used to mine predictive secreted proteins in *F. oxysporum* f. sp. *albedinis* genome.

**Table 2.** Composition of the predicted secretome of F. oxysporum f.sp. albedinis.

| Class | Number of proteins |
| --- | --- |
| Secreted | 1464 |
| Less than 500 amino acids | 1077 |
| No identified pfam domain | 598 |
| Cysteine rich | 550 |
| Cazymes | 386 |
| Repeat containing | 370 |

### A diversified group of necrosis-inducing proteins among candidate effectors of *F. oxysporum* f. sp. *albedinis*

Within fungal secretomes, effectors have been widely investigated for their role in fungal virulence and disease development in plants (Rafiqi et al., 2012, Selin et al., 2016, He et al., 2020). Typically, fungal effector proteins are predicted to function either in the interface between pathogen and host cell structures, or inside host cells, where they manipulate molecular processes in the host in favour of disease development. When they are intercepted by the host immune system, these effectors are called avirulence proteins because their recognition by the host surveillance system limits pathogen virulence and spread (Rafiqi et al., 2012, Vincent et al., 2020, He et al., 2020). Such effectors are under diversification selection to circumvent the host immune surveillance and, therefore, evolve relatively fast, making their prediction challenging. Currently, many thousands of effector candidates have been predicted in genomes of filamentous pathogens and their expression profiles have been analysed, yet for any identified fungal pathogen only a few fungal effectors have been functionally characterised and shown to play a role in fungus-plant interactions (Sánchez-Vallet et al., 2018, Vincent et al., 2020). Nevertheless, certain features have been found to be shared between fungal effectors that make their prediction, albeit inaccurate, possible. Such features include harbouring a signal peptide for secretion outside fungal structures, small size, cysteine content (Saunders et al., 2012, Lu & Edwards, 2016, de Wit, 2016, Jones et al., 2021), a higher sequence diversity than rest of genes (McMullan et al., 2018) or, in rare cases, also harbouring conserved motifs (Liu et al., 2019). To enable uncovering of potential virulence proteins deployed by Foa to invade date palm tissue, we have identified candidate effectors within Foa secretome based on cysteine content (at least 2 counts of predicted disulphide bonds) and using EffectorP 2.0 (Sperschneider et al., 2018) (supplemental data 2). We found that candidate effector proteins account for 30% of the Foa secretome. Among these, we recovered three homologues of necrosis-inducing toxins called Nep1-like proteins (NLPs), which contain the nlp24 peptide with its two conserved regions I (11-aa immunogenic part) and II (the heptapeptide GHRHDWE motif) (Fellbrich et al., 2002). Three other candidate effectors with predicted phytotoxicity and homology to *Cladosporium fulvum* Ecp2 effector (Hce2) (Stergiopoulos et al., 2010) were recovered in Foa secretome (Table 3).

**Table 3.** Potential necrosis-inducing toxins encoded in the *F. oxysporum f*. *sp. albedinis* genome.

| Protein ID | pfam<br>accesssion | pfam names | pfam<br>evalues |
| --- | --- | --- | --- |
| FUN_007374-T1 | PF05630.10 | Necrosis inducing protein (NPP1) | 1.70E-53 |
| FUN_001358-T1 | PF05630.10 | Necrosis inducing protein (NPP1) | 1.10E-56 |
| FUN_000270-T1 | PF05630.10 | Necrosis inducing protein (NPP1) | 8.30E-65 |
| FUN_004711-T1 | PF14856.5 | Pathogen effector putative<br>necrosis-inducing factor (Hce2) | 1.50E-19 |
| FUN_009556-T1 | PF14856.5 | Pathogen effector putative<br>necrosis-inducing factor (Hce2) | 8.00E-30 |
| FUN_007640-T1 | PF14856.5 | Pathogen effector putative<br>necrosis-inducing factor (Hce2) | 5.9E-7 |

Necrosis-inducing secreted proteins have been found in a range of plant-associated microorganisms, including oomycetes, fungi and bacteria and tend to cause necrosis mostly in dicotyledonous plants (Nazar Pour et al., 2020, Gijzen & Nurnberger, 2006). It is unclear whether NLPs identified in Foa genome are causing the necrosis observed in infected monocotyledonous date palm rachises. In Fungi, ascomycetes are known to secrete NLPs that contain six conserved cysteine residues. Interestingly, one of Foa’s predicted NLPs possesses 2 cysteine residues (Table 3), making necrosis-inducing secreted proteins intriguing candidates for experiments to investigate their role in virulence.

Other secreted proteins with putative cytotoxic functions include candidate effectors of predicted S1/P1 Nuclease (FUN_003983-T1) and Ribonucleases T2 (FUN_001364-T1 and FUN_008322-T1) families, that might either act intracellularly as a cytotoxin by scavenging host nucleic acids (Pennington et al., 2019, L. Balabanova, 2012) or extracellularly to inactivate the damage-associated molecular pattern extracellular DNA (Widmer, 2018).

### *F. oxysporum* f. sp. *albedinis* harbours a distinct profile of SIX genes

Fourteen effectors referred to as secreted in xylem (SIX) proteins, some with a proven role in virulence, have been identified in *F. oxysporum* f. sp. *lycopersici* (Rep et al., 2004, Lievens et al., 2009, Thatcher et al., 2012). In tomato and cucurbit-infecting strains, SIX genes were found to be located on small, dispensable accessory chromosomes. Horizontal transfer of these chromosomes from pathogenic lineages of *F. oxysporum* to a non-pathogenic recipient isolate of *F. oxysporum* renders the latter pathogenic on the respective host (van Dam et al., 2017, Ma et al., 2010). Four sequences similar to SIX effectors were recovered from the draft genome of Foa 133 strain. Two sequences are variants of SIX1 and two other sequences are similar to SIX9 and SIX11 (supplemental data 3). Based on this finding, the profile of SIX genes in Foa 133 isolate is distinct from previously identified SIX gene profiles in *formae speciales* of *F. oxysporum*, supporting the hypothesis that each *forma specialis* possesses a unique combination of effectors. However, as only a single Foa strain has so far been sequenced, more population genomics and functional studies will be needed to determine the profile and the function of SIX genes in *F. oxysporum* f. sp. *albedinis*.

### [SG]PC[KR]P effector candidates

We screened the entire predicted secretome of Foa 133 for conserved motifs that have previously been identified in predicted effectors of other filamentous plant pathogens, such as RxLR (Morgan & Kamoun, 2007, Whisson et al., 2007), YxSL[RK] (Levesque et al., 2010), [YFW]xC (Godfrey et al., 2010), [LI]XAR (Yoshida et al., 2009), [RK]CxxCx_12_H (Yoshida et al., 2009), G[I/F/Y][A/L/S/T]R (Catanzariti et al., 2006) and DELD (Zuccaro et al., 2011). While some of these short motifs are frequently found in many Foa secreted proteins, they do not occur at the correct position within Foa secreted proteins and may simply be due to artifacts of random background matches (motif occurrence is included in table S1). On the other hand, 34 putative effectors in Foa’s secretome carry the conserved motif [SG]PC[KR]P immediately following the predicted signal peptide for secretion (supplemental data 4). This motif has, so far, only been detected in Fusarium proteins (Sperschneider et al., 2013). Effector candidates with this motif seem to be rapidly evolving and have been suggested to play a role in pathogenesis (Sperschneider et al., 2015), though functional studies have not as yet been published. [SG]PC[KR]P effector candidates have been identified in the secretomes of *F. graminearum, F. pseudograminearum, F. oxysporum* and *F. solani* secretomes (Sperschneider et al., 2013). In all four species, all proteins carrying this motif also contain cysteines. In Foa’s secretome, [SG]PC[KR]P candidate effectors are predicted to be heavily glycosylated and phosphorylated. They also carry repeats of PAN/apple domains. The PAN/apple domain (PF00024, PF14295) is enriched in the secretomes of oomycete species and is associated with carbohydrate-binding modules, such as cellulose-binding elicitor lectins (CBEL), which also elicit strong host immune responses when infiltrated into host (tobacco) and non-host plants, including *Arabidopsis thaliana* (McGowan et al., 2020, Mesarich et al., 2015, McGowan & Fitzpatrick, 2017, Sejalon-Delmas et al., 1997, Mateos et al., 1997). On the other hand, PAN/apple modules are also involved in protein-protein interactions (Tordai et al., 1999). They are found on Toxoplasma cell surface binding receptors that are involved in cell entry (Gong et al., 2012, Brecht et al., 2001). A noted feature of PAN/apple domain-containing proteins is a conserved pattern of cysteine residues. Seven of the 34 identified Foa [SG]PC[KR]P effector candidates are not predicted to carry any disulphide bond, but are still predicted to carry several CBMs, suggesting a binding activity. Based on homology detection and structure prediction by HMM-HMM comparison (HHpred), [SG]PC[KR]P effector candidates are overall predicted to have a hydrolase activity and to interact with components of the plant cell membrane. This interaction is likely to be mediated through N-terminal noncatalytic CBM domains, whose role could be to bring [SG]PC[KR]P proteins in close proximity to their substrates. In addition, HHpred prediction highlights homology signatures to several membrane-interacting and pore-forming proteins, such as bacterial adhesins, vegetative insecticidal proteins Vip3 and Vip4, viral capsid proteins, and cellulosomal scaffoldin adaptor protein B (table S4).

In Foa 133 genome, [SG]PC[KR]P motif seems to occur exclusively in secreted proteins. Only three non-secreted proteins, FUN_007157-T1, FUN_010458-T1 and FUN_010145-T1 carry [SG]PC[KR]P motif, although not at the right position. Interestingly, one of these hypothetical proteins, FUN_010458-T1, is also predicted to carry two PAN domains. Based on these bioinformatics predictions, we hypothesize that [SG]PC[KR]P effectors could form a machinery of proteins, such as cellulosome or equivalent, in *F. oxyssporum* f. sp. *albedinis*, and perhaps also in other *Fusarium* species. A complex consisting of a variety of different enzymes bound to noncatalytic scaffolding subunits with a role of binding mediators, which can each bind, perhaps specifically hence sequence diversity, to one of the various plant cell surface anchoring proteins. [SG]PC[KR]P proteins will be part of our priority list for mechanistic and functional studies in the date palm dieback pathosystem.

## Conclusions

Palm dieback disease is killing date palms, the cornerstone of life in Saharan oases, and posing a threat to biodiversity as well as to human subsistence and livelihoods in populated oases. We could isolate numerous Foa strains from diseased palms in all five oases we sampled in Morocco (data not shown), indicating a very wide spread of the pathogen within its native range. Palm dieback disease is progressing at an alarming rate, moving West to East across north Africa and reaching as far as the Middle East and Pakistan (Figure 3) (Mirza Hussain Samo, 2020). In an effort to protect date palm cultivars of economic importance and restore it to oases landscapes, millions of genetically diverse palm trees demonstrating different levels of tolerance to the highly destructive palm dieback pandemic have been planted over the last two decades across Morocco’s oases under agroforestry systems (Essarioui, 2020). However, while diversifying host populations has the potential to challenge the pathogen infectivity through variation in host susceptibility, disease progress and spread of infection depends equally on the level of genetic diversity in the pathogen population (Ganz & Ebert, 2010). As such, future research should aim at analysing pathogen strains across infected areas and address more empirical studies. In order to get insights into the mechanisms of infection deployed by Foa to invade palm tissue, we annotated the recently released genome of Foa strain 133 and we analysed its *in silico* secretome, including putative secreted effectors. Two particular sets of secreted proteins, SGPCKRP and SIX effector candidates, are specific to *Fusarium oxysporum* but have their contribution to the pathogen’s virulence as yet uncharacterised. These proteins will be prime candidates for our future studies. Several other effector candidates have predicted cytoplasmic functions and are suggested to be transferred into infected plant cells. These effector candidates are likely to be required more during the initial biotrophic growth of the pathogen than upon its later necrotrophic development. Interestingly, we could isolate Foa strains from many symptom-free palms (data not shown). Fusarium species are likely to have had a long evolutionary history with North African palm trees and may associate with this host in a range of different interactions spanning the spectrum from harmless endophytism to pathogenesis. What caused the outbreak of palm dieback in recent decades is as yet unknown. The increased frequency of severe long and hot drought events in Saharan oases under and ever-changing climate may render the palm dieback pathogen harder to detect and mitigate at the onset of disease development. It is, therefore, timely to address both biochemical and genomics studies in order to elucidate palm dieback disease function and help save date palms, which in turn will help conserve oasitic ecosystems.

**Figure 3.**
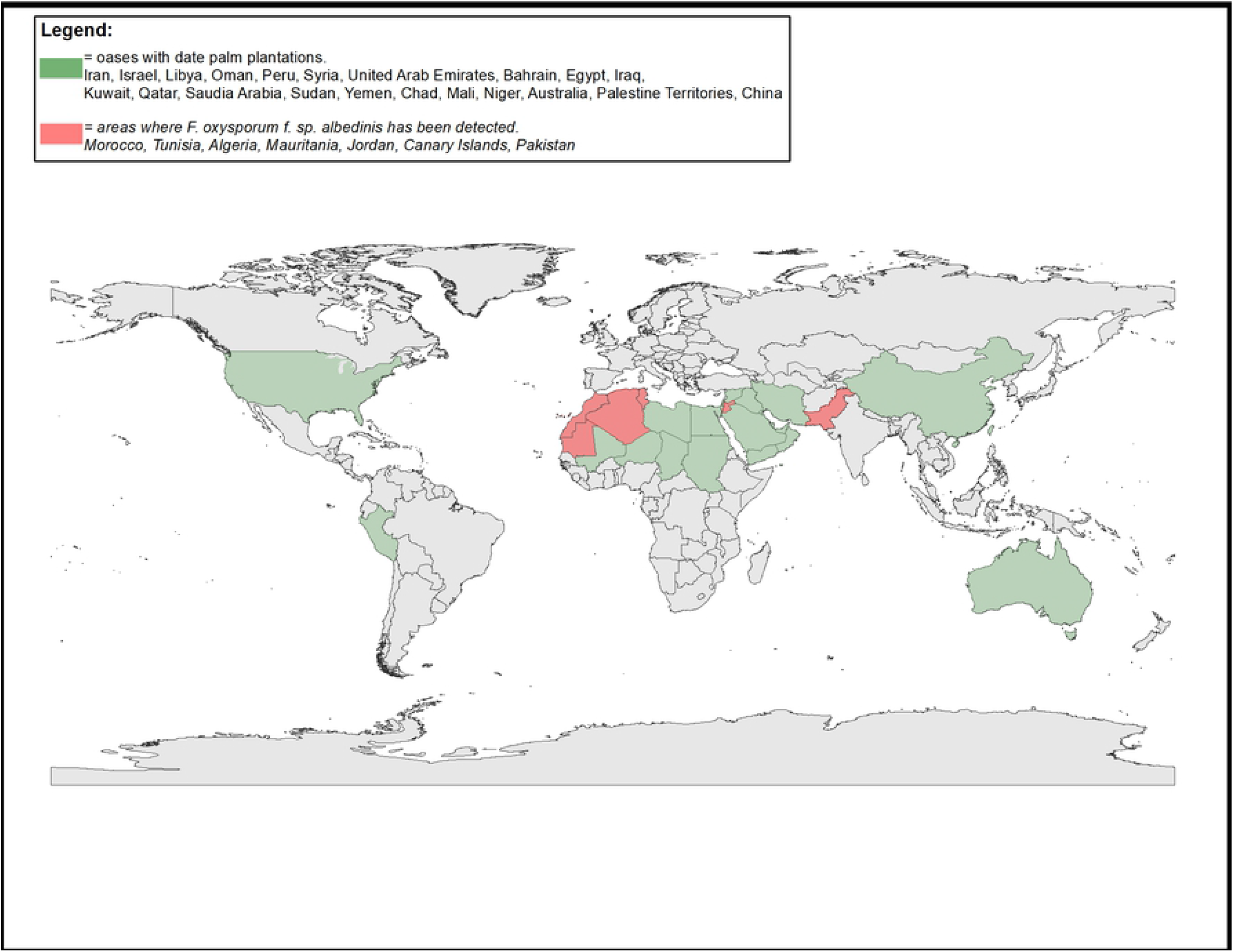
Geographic distribution of the palm dieback disease caused by *F. oxysporum* f. sp. *albedinis*. Map was developed using ArcGIS 10.8 with layer from natural earth dataset. http://www.naturalearthdata.com/

## Materials and Methods

### Annotation of *Fusarium oxysporum* f. sp. *albedinis* strain 133

We used the assembled genome of *Fusarium* oxysporum f. sp. *albedinis* strain 133 (Khayi et al., 2020). Gene prediction and annotation were carried out using funnotate pipeline v1.8.1 (Stajich, 2020), which includes masking, ab initio gene-prediction training, using Augustus and Genmark, with the EST dataset reported to the Ganoderma mycocosm repository, gene prediction, and the assignment of functional annotation to protein-coding gene models.

### Secretome prediction

We used a pipeline described previously (Saunders et al., 2012, Rafiqi et al., 2013) to predict fungal secretomes. Briefly, the pipeline filters a set of proteins that contain signal peptides using SignalP 4. 1f (Nielsen, 2017). This set of predicted secreted proteins were further used in the pipeline to predict transmembrane helices with TMHMM 2.0c (Krogh et al., 2001) and cellular localisation signals with TargetP 1.1b (Emanuelsson et al., 2000). Sequences that contained transmembrane helices or mitochondrial targeting signal were removed from the set of interest. The remaining proteins were annotated with: Hmmer (Zhang et al., 2018) against PfamScan (Finn et al., 2014) for domain information, Targetp (Almagro Armenteros et al., 2019) for subcellular localisation, Predictnls (Cokol et al., 2000) for detection of nuclear localization signals, T-Reks (Jorda & Kajava, 2009) for repeat identification, Disulfinder (database: uniprotkb/swiss-prot) for disulfide bond prediction, MOTIF search for the search of known motifs. The positions of the motifs RxLR, [LI]xAR, [RK]CxxCx{12}H, [YFW]xC, YxSL[RK], G[IFY][ALST]R, DELD and [SG]PC[KR]P are identified with a script based on regular expressions.

### Homology detection and structure prediction

To search for homologues and compare their predicted structure, a workflow that contains the non-modelling steps of the hhpred-website (Zimmermann et al., 2018, Gabler et al., 2020) was implemented. The sequences were scanned with hhblits against uniref30 and pfam to obtain similar sequences, multiple alignments and hmms of the hits and the query sequences were generated, followed by a search of the hmms in pdb with hhsearch. Data were then extracted from hits (see supplemental table S4) and used for the analysis.

## Acknowledgment

This work was supported by University Mohammed VI Polytechnic (UM6P). We thank Dr. Toby Gibson and Dr. Manjeet Kumar for help with effector short linear motif and function predictions. We acknowledge technical assistance by the bioinformatics and systems biology facility of JLU Giessen and the BiGi service center. We thank our field guides in Moroccan oases Dr. M. A. El Houmaizi and Mohamed Bo.

